# mTORC2 loss in oligodendrocyte progenitor cells results in regional hypomyelination in the central nervous system

**DOI:** 10.1101/2022.01.04.474811

**Authors:** Kristin D. Dahl, Hannah A. Hathaway, Adam R. Almeida, Jennifer Bourne, Tanya L. Brown, Lisbet T. Finseth, Teresa L. Wood, Wendy B. Macklin

## Abstract

In the central nervous system (CNS), oligodendrocyte progenitor cells (OPCs) differentiate into mature oligodendrocytes to generate myelin, which is essential for normal nervous system function. OPC differentiation is driven by signaling pathways such as mTOR (Mechanistic Target of Rapamycin), which functions in two distinct complexes: mTOR complex 1 (mTORC1) and mTOR complex 2 (mTORC2), containing Raptor or Rictor respectively. In the current studies, mTORC2 signaling was selectively deleted from OPCs in PDGFRα-Cre X Rictor^fl/fl^ mice. This study examined developmental myelination in male and female mice, comparing the impact of mTORC2 deletion in the corpus callosum and spinal cord. In both corpus callosum and spinal cord, Rictor loss in OPCs resulted in early reduction in myelin RNAs and some myelin proteins. However, these deficits rapidly recovered in spinal cord, where normal myelin abundance and thickness was noted at post-natal day 21 and 1.5 months. By contrast, the losses in corpus callosum resulted in severe hypomyelination, and increased unmyelinated axons. The current studies focus on uniquely altered signaling pathways following mTORC2 loss in developing oligodendrocytes. A major mTORC2 substrate is phospho-Akt-S473, which was significantly reduced throughout development in both corpus callosum and spinal cord at all ages measured, yet this had little impact in spinal cord. Loss of mTORC2 signaling resulted in decreased expression of actin regulators such as gelsolin in corpus callosum, but only minimal loss in spinal cord. The current study establishes a regionally-specific role for mTORC2 signaling in OPCs, particularly in the corpus callosum.

**Significance Statement:** mTORC1 and mTORC2 signaling have differential impact on myelination in the central nervous system. Numerous studies identify a role for mTORC1, but deletion of Rictor (mTORC2 signaling) in late-stage oligodendrocytes had little impact on myelination in the CNS. However, the current studies establish that deletion of mTORC2 signaling from oligodendrocyte progenitor cells results in reduced myelination of brain axons. These studies also establish a regional impact of mTORC2, with little change in spinal cord in these conditional Rictor deletion mice. Importantly, in both, brain and spinal cord, mTORC2 downstream signaling targets were impacted by Rictor deletion. Yet, these signaling changes had little impact on myelination in spinal cord, while they resulted in long term alterations in myelination in brain.

## Introduction

Myelin is a specialized membrane generated by oligodendrocytes in the central nervous system (CNS), which allows salutatory conduction and provides trophic support to axons (Verden & Macklin, 2016). Differentiation of oligodendrocyte precursor cells (OPCs) into myelin-producing oligodendrocytes involves extreme metabolic and morphological changes (Goldman, 1992), including altered myelin gene expression, lipid biosynthesis and cytoskeleton organization, and these changes are driven by selective signaling pathways (Adams et al., 2021).

An important signaling pathway involved in oligodendrocyte differentiation is the mechanistic target of rapamycin (mTOR), a highly conserved serine/threonine kinase that regulates numerous cellular processes (Saxton and Sabatini, 2017). mTOR functions in two distinct complexes: mTOR complex 1 (mTORC1), defined by the presence of Raptor, and mTOR complex 2 (mTORC2), defined by the presence of Rictor (Saxton and Sabatini, 2017, Wood et al., 2013). Numerous studies support an important role for mTOR signaling in OPC differentiation and myelination. Our early *in vivo* studies overexpressing constitutively active Akt in oligodendrocytes established that Akt signaling through mTOR drives oligodendrocyte differentiation and myelination, eventually resulting in hypermyelination (Flores et al., 2008; Narayanan et al., 2009). Pharmacological inhibition of mTOR *in vitro* blocks OPC differentiation (Tyler et al., 2009; Dai et al., 2014), and siRNA knockdown of either Raptor or Rictor decreases myelin protein expression (Tyler et al., 2009). As OPCs differentiate, mTOR regulates myelin protein expression, lipid biosynthesis and cytoskeletal actin dynamics (Bercury et al., 2014; Wahl et al., 2014; Dai et al., 2014; Lebrun-Julien et al., 2014; Mathews and Appel, 2016; Musah et al., 2020).

The two mTOR complexes have different substrates and functions, and the relative contribution of each complex to OPC differentiation and myelination is still poorly understood. Conditional deletion studies establish that mTORC1 (Raptor) or mTORC2 (Rictor) signaling impacts oligodendrocyte differentiation differently, depending on 1) the differentiation stage of oligodendrocytes and 2) the CNS region under investigation. Using the 2’,3’ cyclic nucleotide, 3; phosphodiesterase (CNP) promoter, which expresses Cre late in oligodendrocyte differentiation, mTOR or Raptor conditional deletion results in hypomyelination in spinal cord but not corpus callosum (Bercury et al., 2014; Wahl et al., 2014; Lebrun-Julien et al., 2014). Thus, mTORC1 signaling is required for myelination, although this requirement appears to be unique to specific regions of the CNS. By contrast, ablation of mTORC2 signaling in *CNP-Cre; Rictor^fl/fl^* mice has no impact on myelination in either spinal cord or corpus callosum (Bercury et al., 2014; Lebrun-Julien et al., 2014). Nevertheless, a role for mTORC2 was established by ablating Rictor throughout the oligodendrocyte lineage, i.e., from early developmental stages. Thus, in Olig2-Cre mice, Rictor conditional deletion results in delayed myelination in the corpus callosum (Grier et al., 2017). This suggests that mTORC2 signaling is necessary early in OPC differentiation but that loss of that signaling in CNP-Cre conditional deletion mice is unnecessary or could be compensated for later in development. Grier et al. (2017) did not investigate whether mTORC2 signaling plays a similar role in other regions of the CNS, such as the spinal cord, nor did it investigate the signaling pathways impacted by Rictor deletion.

Olig2-Cre expression in *Olig2-Cre; Rictor^fl/fl^* mice would result in mTORC2 deletion in precursor cells as early as embryonic day 9.5, preceding expression of platelet derived growth factor receptor alpha (PDGFRα) (Zhou and Anderson, 2000). PDGFRα is expressed by OPCs in the CNS, but is downregulated as oligodendrocytes differentiate, making PDGFRα a specific marker of the OPC stage of the oligodendrocyte lineage (Pringle et al., 1992; Pringle and Richardson, 1993). In the current study, we used PDGFRα-Cre mice to delete Rictor, and thereby mTORC2 signaling, selectively in OPCs, specifically impacting differentiation. This resulted in hypomyelination in corpus callosum, but interestingly the impact of Rictor deletion from OPCs was minimal in spinal cord. These data support unique developmental and region-specific requirements of mTOR signaling, and a particular role for mTORC2 signaling in OPC differentiation.

## Materials and Methods

### Mice models

Mice used in this study include PDGFRα-Cre mice (*C57BL/6-Tg(Pdgfrα-cre*) *1Clc/*; JAX Stock No:1013148; RRID:IMSR_JAX:013148; Roesch et al., 2008;) and Rictor^fl/fl^ mice (MMRRC *B6.129S6(SJL)*-*Rictor^tm1.1Mgn^/Mmnc*; RRID:MMRRC_014113-UNC; Shiota et al., 2006). Heterozygous PDGFRα-Cre mice (Cre±) were bred with homozygous Rictor^fl/fl^ mice to obtain *PDGFRα-Cre±;Rictor^fl/+^* mice. These mice were bred with Rictor^fl/fl^ mice to obtain Rictor conditional knockout mice (Rictor cKO; *PDGFRα-Cre±;Rictor^fl/fl^*) and littermate control mice (*PDGFR*-Cre+/+;Rictor^fl/fl^ or *PDGFRα-Cre+/+; Rictor^fl/+^* mice). Both male and female mice were used in all analyses. Genotypes of all mice were determined by PCR using previously published primers and protocols, as well as primers defined by The Jackson Laboratory, oIMR1084 and oIMR1085 (Roesch et al., 2008; Shiota et al., 2006). The Rictor cKO animals (PDGFRα-CRE±; Rictor fl/fl) had a significantly decreased body weight compared to control Cre negative animals. Body weight analysis indicated the mean weight was 7.24g (SEM 0.206; N=22) for control mice and 5.899g (SEM 0.194; N=10) for Rictor cKO mice at P14, using an unpaired t-test with Welch’s correction p=0.000614, df=26.37. At P30 the mean was 16.48g (SEM 0.692, N=7) for control mice and 8.34g (SEM 0.556, N=11) for Rictor cKO mice, using an unpaired t-test with Welch’s correction p=0.0000005; df=13.00. All animal procedures were conducted with the approval of the University of Colorado Institutional Animal Care and Use Committee.

### Electron microscopy

Animals were perfused with modified Karnovsky’s fixative (2% paraformaldehyde/ 2.5% glutaraldehyde) in phosphate buffer, pH 7.4. The brain was removed and immersed in the same fixative overnight. Corpus callosum was isolated from 1 mm coronal slices of brain between -0.94 and -2.18 of bregma (Franklin and Paxinos, 2008). Spinal cords were cut into 500 nm coronal sections through the cervical enlargement. Using a PELCO Biowave Pro tissue processor (Ted Pella), the tissue was rinsed in 100 mM cacodylate buffer and then post-fixed in a reduced osmium mixture consisting of 1% osmium tetroxide and 1.5% potassium ferrocyanide followed by 1% osmium tetroxide alone. Dehydration was carried out in a graded series of acetone (50%, 70%, 90%, 100%) containing 2% uranyl acetate for en bloc staining. Finally, tissue was infiltrated and embedded in Embed 812 (Electron Microscopy Services) and cured for 48 hr at 60°C in an oven. The corpus callosum pieces were oriented such that sections could be cut midline in a sagittal plane. Spinal cord pieces were oriented such that sections could be cut in a transverse plane. Ultrathin sections (65 nm) were mounted on copper grids and viewed at 80 kV on a Tecnai G2 transmission electron microscope (FEI). Electron micrographs of the corpus callosum were imaged near the midline and spinal cord images were obtained from the region of the dorsal columns.

G-ratios were determined using AxonDeepSeg (Zaimi et al., 2018) and manual tracing. A myelin/axon mask was generated and then partial or misidentified objects were manually removed.

### Immunofluorescent and fluorescent staining

Mice were anesthetized with isofluorane followed by intraperitoneal (IP) injection of avertin or Fatal plus. Intracardial perfusion was performed using phosphate buffered saline (PBS) followed by 4% paraformaldehyde (PFA) in PBS with phosphatase inhibitor, 2mM sodium orthovanadate (Na3VO4, CAS: 13721-39-6). Tissues were dissected and then post-fixed in 4% PFA in PBS overnight and transferred to cryoprotect solution (20% glycerol in 0.1 M Sorensen’s buffer or 30% sucrose) for at least 72 hours. Cervical spinal cord and cortex were cut into 30 µm or 20 µm free-floating sections and stored in cryostorage solution (30% ethylene glycol, 30% sucrose, 1% PVP-40 in 0.1 M Sorenson’s buffer). Before staining, antigen retrieval was performed in 10 mM sodium citrate (pH 6) for 5 minutes at 65°C (550W), Pelco Biowave Pro. Sections were permeabilized in 1% Triton-X 100 in Tris buffered saline (TBS) for 10 minutes at room temperature, washed, and then blocked for 1 hour at room temperature with 5% normal donkey serum (NDS) and 0.3% Triton-X 100 in TBS. All other primary antibodies were incubated overnight at 4°C. All secondary antibodies were used at 1:800 concentration and incubated for 45 minutes at room temperature. All primary and secondary antibody solutions were prepared in 3% NDS blocking solution. Sections were rinsed with phosphate buffer and then mounted and cover-slipped with Fluoromount-G (ThermoFisher). For spinal cord analysis, images were acquired in the dorsal column. For corpus callosum, images were acquired at midline in the genu. At least three sections per mouse were imaged.

### Microscopy

Fluorescent images were taken on Nikon A1 Confocal microscope combined with a Ti2-E microscope with a 20x objective. For each experiment all microscopy settings were set the same to maintain consistent and comparable images.

### Immunofluorescent and image analysis

Cell counts using fluorescent images in Figure 2 were done with the automated open source software, Cell profiler (Carpenter et al., 2006). Three to four images for each animal were averaged. The fluorescent intensity was determined using Fiji open-source software (Schindelin et al., 2012). The fluorescent intensity analysis (Figure 3 and 5) workflow was done by generating a 150µm^2^ region of interest (ROI) within the apparent corpus callosum to generate a mean intensity value. This mean intensity was normalized to the average value of the control animals. The F-actin to G-actin ratio was generated using a 150µm^2^ ROI and measuring the F-actin (phalloidin) intensity and the G-actin (DNase I) intensity for the same ROI in the same location, either corpus callosum or cortex. The F-actin intensity measurement was divided by the G-actin intensity measurement.

### Antibodies

Antibodies used for western blot analysis: Total AKT (AKT (pan) (40D4) anti-mouse, Cell Signaling Cat#: 2920S, RRID:AB_1147620, 1:500); CNP (Anti-CNPase antibody produced in rabbit, Sigma-Aldrich SKU:C9743, RRID:AB_1840761, 1:1000); GapDH (GapDH (D16H11) XP Rabbit mAb, Cell Signaling Cat#: 5174S, RRID:AB_10622025, 1:1000); GapDH (GapDH (D4C6R) Mouse mAb, Cell Signaling Cat #: 97166S, RRID:AB_2756824, 1:1000); Gelsolin (Recombinant Anti-Gelsolin antibody [EPR1942], Abcam Cat #: ab109014, RRID:AB_10863643, 1:1000); MAG ((D4G3) XP Rabbit mAb, Cell Signaling Cat #: 9043S, RRID:AB_2665480, 1:1000); MBP (Myelin Basic Protein, Neuromics Cat #: MO22121, RRID:AB_2737143, 1:1000); MOG (Anti-Myelin oligodendrocyte glycoprotein antibody, Abcam Cat #: ab32760, RRID:AB_2145529, 1:1000); Total mTOR (mTOR (7C10) Rabbit mAb, Cell Signaling Cat #: 2983, RRID:AB_2105622, 1:500); pAkt-S473 (Phospho-Akt (Ser473) (D9E) XP Rabbit mAb, Cell Signaling Cat #: 4060S, RRID:AB_2315049, 1:500); pAkt-T308 (Phospho-Akt (Thr308) (D25E6) XP Rabbit mAb, Cell Signaling Cat #: 13038S, RRID:AB_2629447, 1:500); PKCα (D7E6E) Rabbit mAb, Cell Signaling Cat #: 59754S, RRID:AB_2799573, 1:500; PKC alpha/beta II (Phospho-Thr638/641) anti-Rabbit, Cell Signaling Cat #: 9375S, Lot 4; RRID:AB_2284224, 1:1000; PLP, AA3, RRID:AB_2341144, 1:5000; Phospho-mTOR (Ser2481) Antibody, Cell Signaling Cat #: 2974, RRID:AB_2262884, 1:500; Phospho-mTOR (Ser2448) (D9C2) XP Rabbit mAb, Cell Signaling Cat #: 5536P, RRID:AB_10691552, 1:500; Recombinant Anti-Profilin 1 antibody [EPR6304], Abcam ab124904, RRID:AB_10975882, 1:1000; Profilin 2 Antibody, Novus Cat #: 87426, RRID:AB_11007824, 1:250; Phospho-S6 Ribosomal Protein (Ser240/244) Antibody, Cell Signaling Cat #: 2215S, RRID:AB_331682, 1:500; Rictor Antibody anti-Rabbit, Cell Signaling Cat #: 2140, RRID:AB_2179961, 1:500; Tubulin (alpha), Cell Signaling Cat #: 3873S, RRID:AB_2797891, 1:1000. Antibodies used in immunofluorescence staining and actin visualization. CC1 (Anti-APC (Ab-7) Mouse mAb (CC-1), Calbiochem OP80, RRID:AB_2057371, dilution 1:300); MAG (Anti-Myelin Associated Glycoprotein, clone 513 antibody, Millipore mab1567, RRID:AB_2137847, dilution 1:1000); MBP (Myelin Basic Protein, Neuromics Cat #: MO22121, RRID:AB_2737143:, dilution 1:1000); MOG (Myelin oligodendrocyte glycoprotein antibody, Abcam Cat# ab32760, RRID:AB_2145529, dilution 1:1000); Olig2 (Anti-Olig2 antibody, Gift from Charles Stiles, RRID:AB_2336877, dilution 1:200; PLP, AA3, RRID:AB_2341144, 1:100); G-Actin (Deoxyribonuclease I, Alexa Fluor™ 594 Conjugate D12372, Molecular probes, Cat #: D12372); F-actin (Cytoskeleton, Inc. Acti-Stain 488 Phalloidin, Cat #: PHDG1-A)

### Quantitative PCR

Corpus callosum and cervical spinal cord samples at P14 and P30 were dissected, submerged in TRIzol (Invitrogen) and stored at −80°C until extraction. Samples were homogenized and RNA extracted. cDNA was generated from 1 µg of RNA using iScript^TM^ Reverse Transcription Supermix for RT-qPCR (Bio-Rad). Real-time quantitative PCR was performed on a StepOnePlus real-time PCT Machine (Applied Biosystems). Taqman Universal PCR Master Mix (Thermo Fisher) was used with Taqman probes (Thermo Fisher) *gapdh* (Ref 4352661), *rictor* (Mm01307318_m1), *cnp* (Mm01306641), *mog* (Mm00447824_m1), *mbp* (Mm01266402_m1), *plp1* (Mm01297210_m1), and *mag* (Mm00487538_m1). All experiments followed the MIQE Guidelines for qPCR (Bustin et al., 2009).

### Western blots

The corpus callosum and cervical spinal cord of P7, P14, and P30 animals was dissected, snap frozen in liquid nitrogen, and stored at −80°C until lysis. The tissue was lysed in a glass homogenizer in cold RIPA buffer (Sigma-Aldrich) with a phosphatase inhibitor cocktail (Calbiochem; Millipore) and protease inhibitor tablet (complete-mini; Roche). The crude lysates were centrifuged at 10,000 × *g* for 10 min at 4°C. The supernatants were then collected, and the protein concentration was determined using the BCA protein Assay Kit (Thermo Fisher). The lysates were then resolved using 4-20% SDS-PAGE gradient gels and transferred to PVDF membranes. The membranes were blocked with 5% bovine serum albumin in TBS and then incubated with primary antibodies overnight at 4°C. Immunodetection was performed using IRDye infrared secondary antibodies (LI-COR). All blots were scanned and quantified using an Odyssey Infrared imager (LI-COR) and Image Studio Lite software (Ver 5.2).

### Experimental Design and Statistical Analysis

All statistical analyses were done using GraphPad Prism version 9.3.1 for Windows, GraphPad Software, San Diego, California USA, www.graphpad.com. When testing significance between the conditional knockout and control sample, we used unpaired t test with Welch’s correction to account for variability in the standard deviation. In Figure 7 we analyzed total axon diameter by g-ratio and performed a simple linear regression for the graphs. Figure 7 also included a histogram analysis of the relative frequency, in which we used frequency distribution tabulated as relative frequency in percentages with a bin width of 0.5. In Figure 8 when testing the significance between cortex staining and corpus callosum staining a paired t test was used to account for batch variability. Statistical significance was defined as p < 0.05. When error bars are shown they are reported as the mean value ± standard mean error.

### Data access

Data is made available on https://osf.io/

## Results

### CNS region-specific myelin deficiency in PDGFRα-Cre; Rictor^fl/fl^ mice

To investigate the role of mTORC2 signaling in OPC myelination, we crossed PDGFRα-Cre mice with Rictor^fl/fl^ mice to generate *PDGFR*α*-Cre; Rictor^fl/fl^* mice (Rictor cKO mice). FluoroMyelin staining of brain and spinal cord from postnatal day 30 (P30) Rictor cKO mice and Rictor^fl/fl^ littermate controls demonstrated dramatic reduction of myelin in brain, with little reduction in spinal cord (Figure 1A, 1B), which suggests a region-specific role of mTORC2 signaling in PDGFRα-expressing OPCs.

### Rictor loss reduced oligodendrocyte differentiation in corpus callosum but not spinal cord

Since reduced mTORC2 signaling in OPCs apparently reduced myelin in corpus callosum but not spinal cord (Figure 1), it was important to test whether OPC differentiation was affected in corpus callosum or spinal cord. We quantified the number of total oligodendrocytes and the number of differentiating cells in Rictor cKO mice by quantifying the total number of Olig2+ oligodendrocyte lineage cells; differentiating cells were quantified by determining the relative percentage of CC1+ cells among total Olig2+ oligodendrocytes. In corpus callosum, there was a statistically significant decrease in the total number of Olig2+ cells in Rictor cKO mice, which was further decreased by P30 (Figure 2C). The percent of those cells that differentiated to CC1+ cells was also reduced in corpus callosum at P14, but not by P30 (Figure 2D). Thus, the relative number of oligodendrocyte lineage cells that were differentiating to CC1-positive cells was initially only about 50% of the low number of total Olig2-positive cells at P14. By P30, the relative number of differentiating cells was comparable to control, suggesting differentiation was delayed. However, there were far fewer oligodendrocyte lineage cells by P30 in corpus callosum, and given the dramatic reduction in total Olig2+ cells at P30, the overall reduction in CC1-positive cells in corpus callosum was large. As expected, in spinal cord where most measures of myelination were similar to control, there was no change in the total number of Olig2 cells at either age, or of the percentage of CC1+ cells among oligodendrocytes in Rictor cKO compared to control (Figure 2E,F).

**Figure 1:**
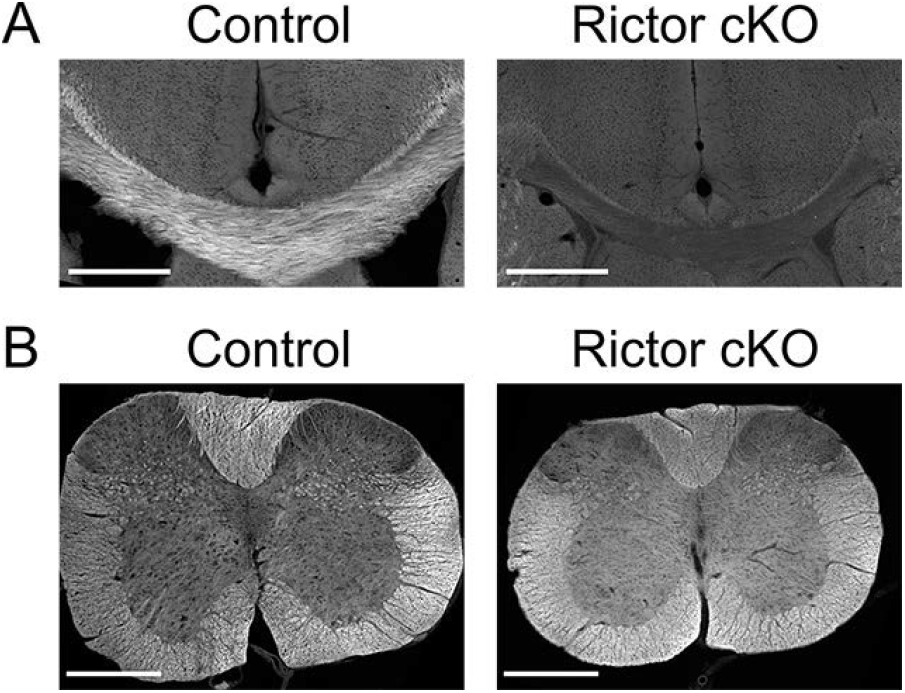
Rictor conditional knockout (cKO) in OPCs results in hypomyelination in the corpus callosum and not in the spinal cord. Brain or spinal cord sections (P30) were stained with FluoroMyelin. (A) Corpus callosum in *PDGFRα-Cre; Rictor^fl/fl^* mice had reduced FluoroMyelin signal compared to the corpus callosum in control mice. Scale bar 500µm. (B) *PDGFRα-Cre; Rictor^fl/fl^* spinal cord had no apparent reduction of FluoroMyelin signal compared to control. Scale bar 500µm.

**Figure 2:**
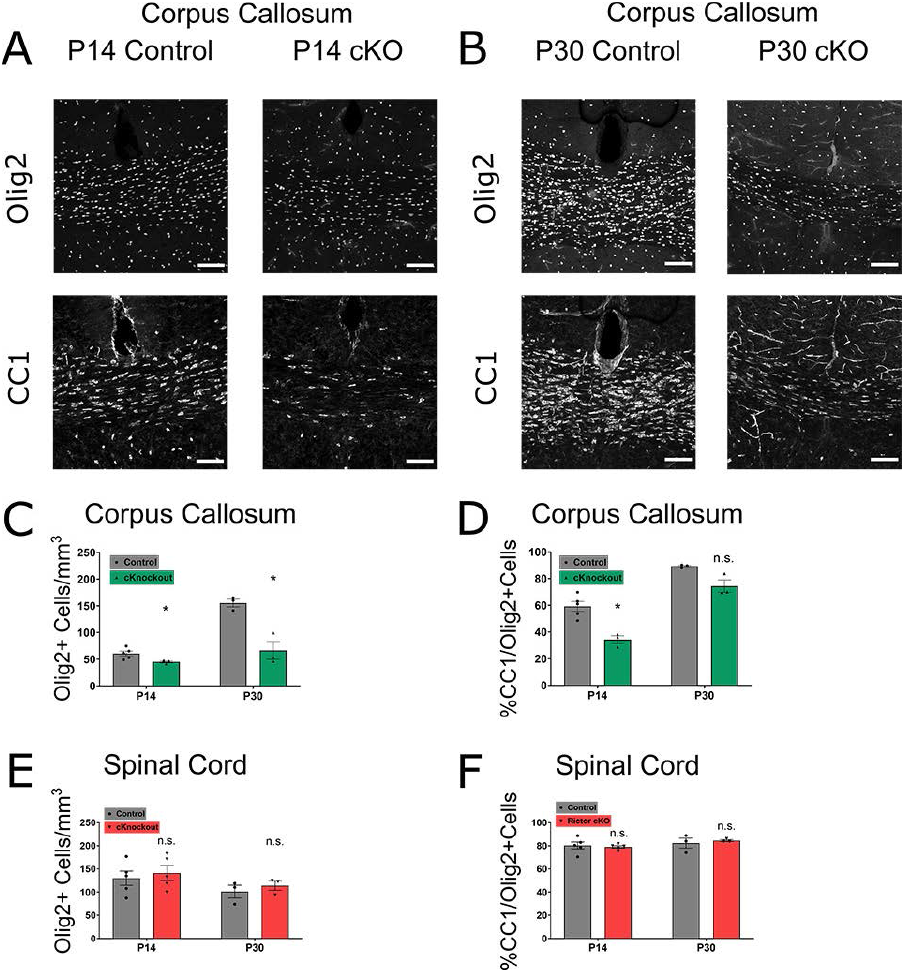
Loss of mTORC2 signaling results in decreased differentiation of Olig2+ cells in the corpus callosum but not the spinal cord at P14 and decreased overall Olig2+ cells at P30. **(A-B)** Olig2 and CC1 staining in corpus callosum at P14 and P30 in control compared to Rictor conditional knockout (cKO). Scale bar 100µm. (**C**) Corpus callosum quantification of Olig2 positive cells per mm^3^. P14 (control n=5, SEM ±4.568; mean=60.65 cells/mm^3^; cKO n=3, SEM±1.952; mean=45.05 cells/mm^3^, p=0.0240, df=5.244); P30 (control n=3, SEM±7.278; mean=155.4 cells/mm^3^; cKO n=3, SEM ±16.63; mean=66.36 cells/mm^3^, p=0.0200, df=2.739) (**D**) Percent of CC1-Olig2 double positive cells to total Olig2 positive cells in the corpus callosum. P14 (control n=5, SEM±3.712; mean=59.2%cells; cKO n=3, SEM±2.911; mean=34.3%cells, p=0.0019, df=5.940); P30 (control n=3, SEM±0.689; mean=89.3%cells; cKO n=3, SEM±4.648; mean=74.4%cells, p=0.0813, df=2.088) (**E**) Spinal cord quantification of Olig2 positive cells per mm^3^. P14 (control n=5, SEM±15.33, mean=129.9 cells/mm^3^; cKO n=5, SEM±15.71; mean=140.9 cells/mm^3^, p=0.6288, df=7.995); P30(control n=3, SEM±13.74, mean=101.1 cells/mm^3^; cKO n=3, SEM±9.984; mean=113.9 cells/mm^3^, p=0.4961, df=3.652) (**F**) Percent of CC1-Olig2 double positive cells to total Olig2 positive cells in the spinal cord. P14 (control n=5, SEM±2.983, mean=80.20%cells; cKO n=5, SEM±1.079; mean=78.96%cells, p=0.7114, df=5.030); P30(control n=3, SEM±4.344, mean=82.22%cells; cKO n=3, SEM±0.852; mean=84.40, p=0.6689, df=2.154).

### Decreased myelin gene expression following Rictor deletion from OPCs

We hypothesized that the loss of mTORC2 signaling would result in decreased myelin protein expression in the corpus callosum consistent with the decreased FluoroMyelin staining (Figure 1) and reduced oligodendrocyte number and differentiation (Figure 2). We therefore investigated myelin protein and mRNA expression in corpus callosum (Figure 3). All myelin specific proteins studied, i.e., proteolipid (PLP), CNP, myelin associated glycoprotein (MAG), myelin-oligodendrocyte glycoprotein (MOG) and myelin basic protein (MBP) were significantly decreased in Rictor cKO mice compared to control at P14, by western blot analysis (Figure 3). Immunofluorescence images of each myelin protein stain were quantified at P14, and these show a statistically significant decrease in mean intensity in all staining except CNP, which had a higher background signal (Figure 3). CNP, MAG, MOG and MBP proteins were still reduced at P30 (Figure 3B’-E’), although PLP protein level had recovered (Figure 3A’). Thus, the impact of the loss of mTORC2 signaling in oligodendrocytes extended into early adulthood, with some myelin specific proteins, such as PLP, recovering with age.

**Figure 3:**
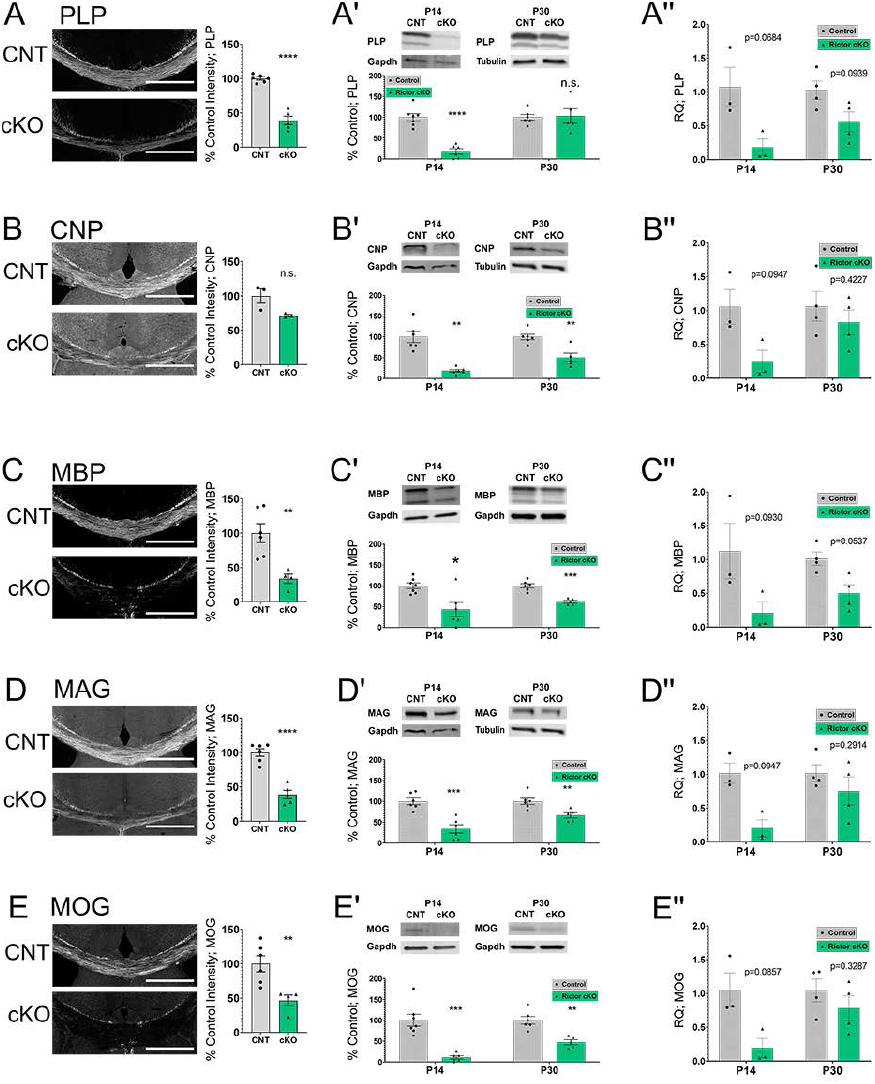
Myelin specific proteins are decreased in the Rictor cKO (cKO) compared to control (CNT) in the corpus callosum through development. (A) Immunofluorescent (IF) images at postnatal day 14 (P14) of proteolipid protein (PLP) in the genu of the corpus callosum. PLP-P14 (control [CNT] n=6, SEM±2.25; cKO n=5, SEM±6.12; cKO mean=39.00%, p=0.0002, df=5.075). **(A’ and A”) PLP protein and RNA expression in corpus callosum at P14 and P30. (A’)** Western blot (WB): PLP-P14 (CNT n=7, SEM±8.49; cKO n=6, SEM±6.44; cKO mean=17.99%, p=0.00001, df=10.66), P30 (CNT n=6, SEM±6.90; cKO n=5, SEM±17.3; cKO mean=103.2%, p=0.8688, df=5.267); (**A’’**) qPCR shown as RQ: P14 (CNT n=3, SEM±0.29; CNT mean=1.07, cKO n=3, SEM±0.12; cKO mean=0.18, p=0.0684, df=2.567); P30(CNT n=4, SEM±0.14; CNT mean=1.03, cKO n=4, SEM±0.15; cKO mean=0.56, p=0.0939, df=4.176) **(B) IF images at P14 2′,3′-Cyclic nucleotide 3′-phosphodiesterase (CNP) in the corpus callosum**. CNP-P14 (CNT n=3, SEM ±10.42; cKO n=3, SEM ±1.74; cKO mean=71.58%, p=0.1085, df=2.111). (**B’ and B”) CNP protein and RNA expression in corpus callosum at P14 and P30. (B’)** WB: CNP-P14 (CNT n=6, SEM±13.24; cKO n=6, SEM±3.17; cKO mean=18.59%, p=0.0013, df=5.573), P30 (CNT n=6, SEM±6.52; cKO n=5, SEM±10.60; cKO mean=50.48%, p=0.0056, df=6.822). (B’’) qPCR: P14 (CNT n=3, SEM±0.26; CNT mean=1.06, cKO n=3, SEM±0.16; cKO mean=0.25, p=0.0947, df=2.466); P30 (CNT n=4, SEM±0.22; CNT mean=1.06, cKO n=4, SEM±0.18; cKO mean=0.83, p=0.4227, df=5.758) **(C) IF images at P14 Myelin Basic Protein (MBP) in the corpus callosum.** MBP-P14 (CNT n=6, SEM ±5.51; cKO n=5, SEM ±6.13; cKO mean=38.98%, p=0.00005, df=8.591). **(C’ and C”) MBP protein and RNA expression in corpus callosum at P14 and P30. (C’)** WB: MBP-P14 (CNT n=7, SEM±6.78; cKO n=6, SEM±16.91; cKO mean=43.42%, p=0.0185, df=6.593), P30(CNT n=6, SEM±5.02; cKO n=5, SEM±3.34; cKO mean=61.79%, p=0.0002, df=8.359). (C’’) qPCR: P14 (CNT n=3, SEM±0.41; CNT mean=1.13, cKO n=3, SEM±0.16; cKO mean=0.21, p=0.0930, df=2.685), P30 (CNT n=4, SEM±0.10; CNT mean=1.01, cKO n=4, SEM±0.13; cKO mean=0.5, p=0.0537, df=3.735) **(D) IF images at P14 Myelin-associated glycoprotein (MAG) in the corpus callosum.** MAG-P14 (CNT n=6, SEM ±13.78; cKO n=4, SEM ±6.66; cKO mean=33.70%, p=0.0035, df=6.972). (D’ and D”) MAG protein and RNA expression in corpus callosum at P14 and P30. (**D’**) WB: MAG-P14 (CNT n=6, SEM±7.96; cKO n=6, SEM±10.02; cKO mean=33.65%, p=0.0005, df=9.514), P30(CNT n=6, SEM±7.47; cKO n=5, SEM±6.21; cKO mean=66.69%, p=0.0076, df=8.954); (D’’) qPCR: P14 (CNT n=3, SEM±0.15; CNT mean=1.02, cKO n=3, SEM±0.13; cKO mean=0.20, p=0.0930, df=2.685), P30(CNT n=4, SEM±0.12; CNT mean=1.02, cKO n=4, SEM±0.21; cKO mean=0.75, p=0.2914, df=3.678) **(E) IF images at P14 Myelin oligodendrocyte glycoprotein (MOG) in the corpus callosum.** MOG-P14 (CNT n=6, SEM ±11.73; cKO n=4, SEM ±8.02; cKO mean=46.47%, p=0.0056, df=7.893). (**E’ and E”) MOG protein and RNA expression in corpus callosum at P14 and P30. (E’)** WB: MOG-P14 (CNT n=7, SEM±14.26; cKO n=6, SEM±3.71; cKO mean=11.05%, p=0.0006, df=6.804), P30(CNT n=6, SEM±9.06; cKO n=5, SEM±5.99; cKO mean=47.29%, p=0.0011, df=8.338); (**E**’’) qPCR: P14 (CNT n=3, SEM±0.25; CNT mean=1.05, cKO n=3, SEM±0.14; cKO mean=0.19, p=0.0857, df=2.357), P30(CNT n=4, SEM±0.17; CNT mean=1.05, cKO n=4, SEM±0.18; cKO mean=0.79, p=0.3287, df=5.450). CNT mean set to 100% based on averaging of actual numerical values for IF and WB. Values are displayed as ±SEM. Unpaired t-test with Welch’s correction. Scale bar 500µm.

Because mTOR can regulate both protein translation and RNA transcription, it was important to establish whether the decreased myelin proteins in the Rictor cKO resulted from reduced protein translation or whether myelin RNAS were reduced. qPCR of Rictor cKO and control tissue established that myelin mRNAs were reduced in the Rictor cKO mice compared to control at P14. However, given sample variability we used a more stringent statistical test (see methods section), and the differences were not significant (Figure 3 A”-E’’). At P30, myelin mRNA levels approached control levels in the Rictor cKO corpus callosum. Thus, the loss of myelin proteins in the Rictor cKO corpus callosum likely resulted from decreased mRNA abundance, at least at P14, but there is recovery with time. Together these data demonstrate that mTORC2 signaling positively impacts oligodendrocyte differentiation and consequently myelin gene expression in the corpus callosum during development.

#### Decreased mTORC2 signaling in Rictor cKO mice

It was important to establish if Rictor cKO mice had reduced mTORC2 signaling in regions with reduced myelin. Importantly, Rictor protein expression was significantly reduced at P7, P14, and P30 in Rictor cKO corpus callosum (Figure 4A), establishing effective deletion of Rictor. Signaling through the mTORC2 complex (Figure 4B) was investigated to determine whether the deletion of Rictor resulted in a loss of mTORC2 signaling. We analyzed the phosphorylation of mTORC2 specific substrates Akt-S473 and pPKCα/β in Rictor cKO and control corpus callosum, and both were significantly decreased in Rictor cKO mice at P7, P14 and P30 (Figure 4C,D). We also measured the total protein levels of PKCα and found that the levels of total PKCα were significantly decreased at all ages, which may account for some but likely not all of the decreased pPKCα/β (Figure 4E). Thus, Rictor cKO mice had long-term deletion of Rictor protein and of mTORC2 signaling in corpus callosum through at least age P30.

**Figure 4:**
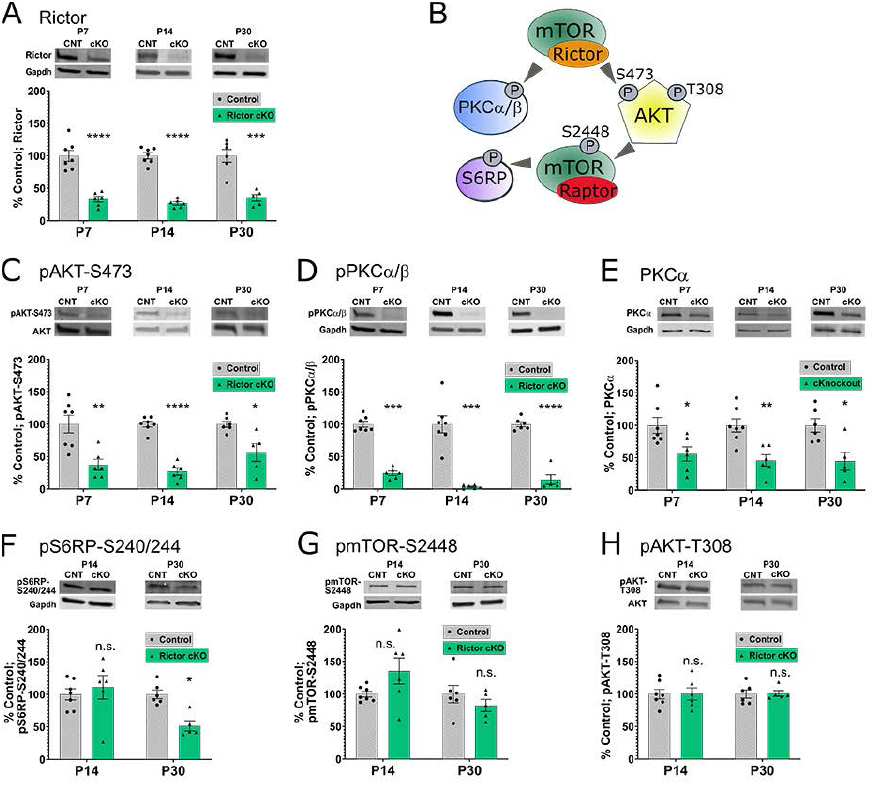
Rictor protein expression and mTORC2 signaling is reduced in the corpus callosum of Rictor conditional through development. **(A) Rictor protein expression normalized to GapDH in Rictor cKO (cKO) compared to control (CNT) at P7, P14 and P30.** P7 (CNT n=7, SEM±8.44; cKO n=6, mean=33.12%, SEM±4.35, p=0.00007; df=8.857). P14 (CNT n=7, SEM±4.66, cKO n=6, mean=26.76%, SEM±2.53; p=0.0000002, df=9.106). P30 (CNT n=6, SEM ±9.83; cKO n=5, mean=35.14%, SEM ±5.20; p=0.0005, df=7.460). **(B) Model of mTORC2 interaction with substrates in the mTOR pathway. (C-H) Western blots of mTORC1 and mTORC2 substrates. (C-D) Western blot of phosphorylated mTORC2 substrates. (C) pAKT:** P7 (CNT n=7, SEM±13.67; cKO n=6, mean=37.32%, SEM±8.45; p=0.0031, df=9.753). P14 (CNT n=7, SEM±4.21; cKO n=6, mean=26.67%, SEM±5.50; p=0.000001, df=9.780). P30 (CNT n=6, SEM±4.67; cKO n=5, mean=56.27%, SEM±13.61; p=0.0292, df=4.944). **(D) pPKCα/β**: P7 (CNT n=7, SEM±4.43; cKO n=6, mean=24.55%, SEM±3.29; p= 0.00000005, df=10.58l). P14 (CNT n=7, SEM±13.23; cKO n=6, mean=3.46%, SEM±1.09; p=0.0003, df=6.081). P30 (CNT n=6, SEM±3.81; cKO n=5, mean=14.63%, SEM±7.54; p=0.00005, df=5.994). **(E) total PKCα**: P7 (CNT n=7, SEM±12.19; cKO n=6, mean=55.97%, SEM±10.90; p=0.0209, df=11.00). P14 (CNT n=7, SEM±10.19; cKO n=6, mean=45.81%, SEM±8.860; p=0.0020, df=10.97). P30 (CNT n=6, SEM±10.45; cKO n=5, mean=44.95%, SEM±13.81; p=0.0134, df=7.834). **(F-H) Western blot of phosphorylated non-mTORC2 substrates. (F) pS6RP: P14** (CNT n=7, SEM±8.43; cKO n=6, mean=110.80%, SEM±18.14; p=0.6066, df=7.109). P30 (CNT n=6, SEM±6.57; cKO n=5, mean=51.50%, SEM±7.73; p=0.0012, df=8.373). **(G) pmTOR-S2448**: P14 (CNT n=7, SEM±4.37; cKO n=6, mean=135.4%, SEM±20.10; p=0.1403, df=5.473). P30 (CNT n=6, SEM±13.19; cKO n=5, mean=82.47%, SEM±9.38; p=0.3083, df=8.591). **(H) pAKT-T308:** P14 (CNT n=7, SEM±7.24; cKO n=6, mean=100.2%, SEM±9.28; p=0.9859, df=9.888). P30 (CNT n=6, SEM±6.29; cKO n=5, mean=101.1%, SEM±4.37; p=0.8891, df=8.517).

Because mTORC2 functions within the larger mTOR signaling pathway, we investigated whether other parts of that pathway were impacted by the loss of mTORC2 signaling. No significant change in the total protein levels of Akt, mTOR, or S6RP were noted, although at P30 there was a significant 13% decrease in total Akt (data not shown). Phosphorylation at pAkt-T308 and pmTOR-S2448 were unaffected at P14 and P30 (Figure 4F,G). Thus, initial activation of Akt from upstream signaling (pAkt-T308) was normal, as was mTOR phosphorylation normally regulated by mTORC1 (Copp et al., 2009). Downstream signaling through mTORC1 was initially unaffected, i.e., there was no significant reduction of pS6RP at P14. Interestingly, however, at P30 there was a significant decrease in pS6RP in Rictor cKO corpus callosum compared to control. Together these data indicate that the loss of mTORC2 signaling did not initially impact major mTORC1 signaling, although the loss of pS6RP by P30 may result from the overall impact of mTORC2 loss on OPC differentiation in corpus callosum feeding back to reduce signaling regulating overall myelination.

#### Impact of mTORC2 loss in spinal cord OPCs

Our initial FluoroMyelin staining showed that Rictor loss in OPCs had no noticeable effect on spinal cord myelination (Figure 1), or oligodendrocyte differentiation (figure 2) suggesting region-specific mTORC2 signaling. It was important then to assess myelin gene expression and mTORC2 signaling in Rictor cKO spinal cord. When myelin protein expression was quantified in Rictor cKO spinal cord, PLP and MAG were not significantly changed compared to control at either P14 or P30 (Figure 5). By contrast, CNP and MBP protein levels were initially significantly decreased in Rictor cKO spinal cord relative to P14 controls, and MOG protein was somewhat non-significantly reduced at P14. By P30, however, CNP and MBP were no longer decreased, but interestingly, MOG protein was significantly decreased in Rictor cKO spinal cord compared to control at P30. While the myelin protein levels at P14 were somewhat reduced, the RNAs for all these proteins were dramatically reduced at P14 in Rictor cKO spinal cord relative to controls. However, these myelin RNAs had all recovered to control levels by P30 in this tissue.

**Figure 5:**
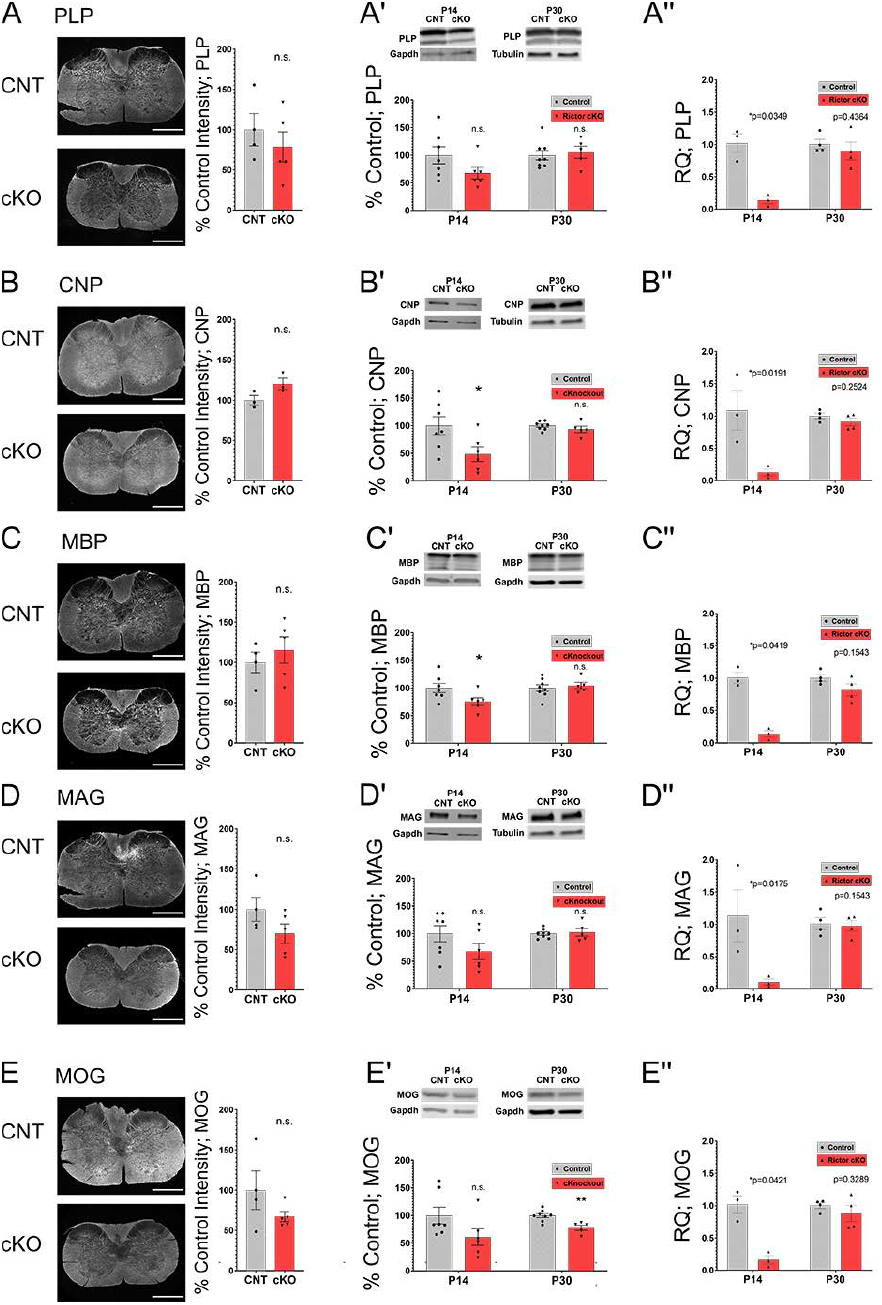
Myelin specific proteins are similar in the cKO compared to CNT in the spinal cord through development. (A) Immunofluorescent (IF) images at P14 of proteolipid protein (PLP) in the spinal cord. PLP-P14 (CNT n=4, SEM±20.02; cKO n=5, SEM±18.67; cKO mean=78.51%, p=0.4592, df=6.691). (**A’ and A”) PLP protein and RNA expression in spinal cord at P14 and P30. (A’**) Western blot (WB): PLP-P14 (CNT n=7, SEM±15.49; cKO n=6, SEM±10.63; cKO mean=67.65%, p=0.1151, df=10.25), P30(CNT n=8, SEM±8.58; cKO n=5, SEM±11.04; cKO mean=105.4%, p=0.7114, df=8.514). (**A**’’) qPCR as shown by RQ: P14(CNT n=3, SEM±0.14; CNT mean=1.02, cKO n=3, SEM±0.05; cKO mean=0.14, p=0.0349, df=2.405), P30(CNT n=4, SEM±0.07; CNT mean=1.01, cKO n=4, SEM±0.14; cKO mean=0.90, p=0.4364, df=4.260) **(B) IF images at P14 2′,3′-Cyclic nucleotide 3′-phosphodiesterase (CNP) in the spinal cord.** CNP-P14 (CNT n=3, SEM±5.87; cKO n=3, SEM±7.08; cKO mean=120.2%, p=0.0951, df=3.869). **(B’ and B”) CNP protein and RNA expression in spinal cord at P14 and P30. (B’)** Western blot: CNP-P14 (CNT n=7, SEM±16.36; cKO n=6, SEM±13.22; cKO mean=48.79%, p=0.0334, df=10.84); P30(CNT n=8, SEM±2.68; cKO n=5, SEM±5.71; cKO mean=93.38%, p=0.3354, df=5.792). (**B**’’) qPCR: P14(CNT n=3, SEM±0.30; CNT mean=1.09, cKO n=3, SEM±0.05; cKO mean=0.13, p=0.0191, df=3.405), P30(CNT n=4, SEM±0.04; CNT mean=1.00, cKO n=4, SEM±0.06; cKO mean=0.91, p=0.2524, df=5.129) **(C) IF images at P14 Myelin Basic Protein (MBP) in the spinal cord.** MBP-P14 (CNT n=4, SEM±12.66; cKO n=5, SEM±16.40; cKO mean=115.3%, p=0.4837, df=6.814). (**C’ and C”) MBP protein and RNA expression in spinal cord at P14 and P30. (C’**) Western blot: MBP-P14 (CNT n=7, SEM±8.43; cKO n=6, SEM±6.81; cKO mean=75.57%, p=0.0459, df=10.85), P30 (CNT n=8, SEM±5.92; cKO n=5, SEM±5.81; cKO mean=104.4%, p=0.6068, df=10.29). (C’’) qPCR: P14(CNT n=3, SEM±0.09; CNT mean=1.01, cKO n=3, SEM±0.05; cKO mean=0.14, p=0.0419, df=2.122), P30(CNT n=4, SEM±0.05; CNT mean=1.00, cKO n=4, SEM±0.09; cKO mean=0.82, p=0.1543, df=4.131) **(D) IF images at P14 Myelin-associated glycoprotein (MAG) in the spinal cord**. MAG-P14 (CNT n=4, SEM±14.95; cKO n=5, SEM±11.83; cKO mean=69.99%, p=0.1654, df=6.132). (**D’ and D”) MAG protein and RNA expression in spinal cord at P14 and P30. (D**’) MAG-P14 (CNT n=7, SEM±14.70; cKO n=6, SEM±14.71; cKO mean=68.04%, p=0.1527, df=10.91), P30 (CNT n=8, SEM±2.91; cKO n=5, SEM±7.31; cKO mean=102.7%, p=0.7444, df=5.291). (D’’) qPCR: P14(CNT n=3, SEM±0.40; CNT mean=1.13, cKO n=3, SEM±0.05; cKO mean=0.11, p=0.0175, df=3.647), P30(CNT n=4, SEM±0.09; CNT mean=1.01, cKO n=4, SEM±0.08; cKO mean=0.98, p=0.8009, df=5.999) **(E) IF images at P14 Myelin oligodendrocyte glycoprotein (MOG) in the spinal cord**. MOG-P14 (CNT n=4, SEM±23.90; cKO n=5, SEM±6.17; cKO mean=67.11%, p=0.2650, df=3.402). (**E’ and E”) MOG protein and RNA expression in spinal cord at P14 and P30. (E**’) Western blot: MOG-P14 (CNT n=7, SEM±15.00; cKO n=6, SEM±14.94; cKO mean=61.07%, p=0.0399, df=10.92), P30(CNT n=8, SEM±3.62; cKO n=5, SEM±4.62; cKO mean=78.03%, p=0.0056, df=8.565). (E’’) qPCR: P14(CNT n=3, SEM±0.14; CNT mean=1.02, cKO n=3, SEM±0.07; cKO mean=0.16, p=0.0421, df=2.349); P30(CNT n=4, SEM±0.04; CNT mean=1.00, cKO n=4, SEM±0.12; cKO mean=0.88, p=0.3289, df=6.000). CNT mean set to 100% based on averaging of actual numerical values for IF and WB. Values are displayed as ±SEM. Unpaired t-test with Welch’s correction. Scale bar 500µm.

Given the apparently differential impact of Rictor loss in developing OPCs in spinal cord relative to corpus callosum, we investigated mTORC2 signaling in developing spinal cord. As in corpus callosum, there was a significant decrease in Rictor protein levels in the spinal cord of Rictor cKO mice (Figure 6A) at P7, P14 and P30. There was also a decrease in the mTORC2 signaling substrate pAkt (S473) (Figure 6B). Unexpectedly, we were surprised to find that neither control nor Rictor cKO spinal cord had detectable signal for phosphorylation of the mTORC2 substrate pPKCα/β (Figure 6C). To confirm its absence in spinal cord, we compared pPKCα/β expression in corpus callosum and spinal cord from P7 to P30 (Figure 6C). At no time during development was pPKCα/β detected in control or Rictor cKO spinal cord, while significant expression was noted in control corpus callosum at all ages, which was reduced in Rictor cKO corpus callosum. Total levels of PKCa were measured and determined to be significantly decreased in the Rictor cKO compared to control, however as we did not observe any pPKCα/β in either the control or cKO spinal cord this decrease in total protein would not likely contribute to decreased signaling (Figure 6D). As in Rictor cKO corpus callosum, there was no significant change in mTORC1 specific substrates in the spinal cord (Figure 6E-G), including pS6RP, which was reduced in older Rictor cKO corpus callosum, and which we hypothesized may result from the overall reduction in oligodendrocyte differentiation/myelination in corpus callosum (Figure 2, Figure 4E). Total levels of Akt, mTOR and S6 were measured; only total mTOR at P30 was significantly decreased by 38% (data not shown). These data confirm the loss of Rictor protein and of classical mTORC2 signaling in the spinal cord of Rictor cKO mice, but despite these losses, no significant myelin loss was noted during Rictor cKO spinal cord development (Figure 5). These data further demonstrate significant differences in the mTORC2 signaling requirements for corpus callosum and spinal cord oligodendrocytes.

**Figure 6:**
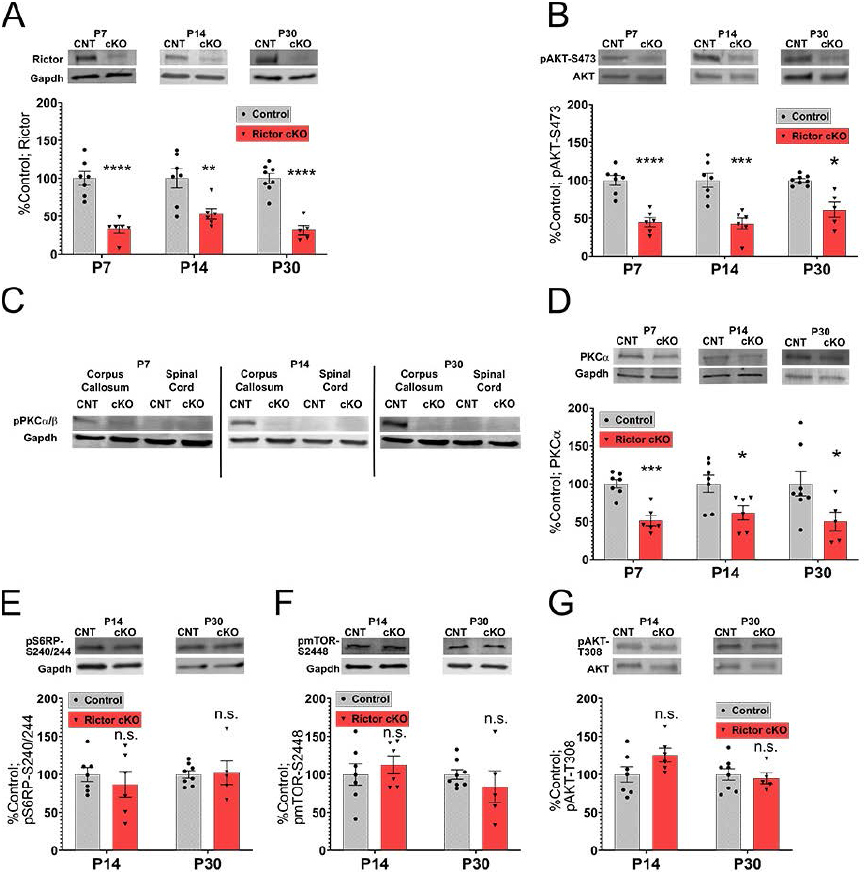
Rictor protein expression and mTORC2 signaling is reduced in the spinal cord of Rictor conditional knockout through development. (A) Rictor protein expression normalized to GapDH in Rictor cKO (cKO) compared to control (CNT) at P7, P14 and P30. P7 (CNT n=7, SEM±9.03; cKO n=6, mean=33.08%, SEM±5.50, p=0.0001; df=9.678). P14 (CNT n=7, SEM±12.52, cKO n=6, mean=53.15, SEM±6.86; p=0.0093, df=9.153). P30 (CNT n=8, SEM±6.37; cKO n=5, mean=31.75%, SEM±6.25; p=0.00001, df=10.28. (B) pAKT-S473 normalized to total AKT in cKO compared to CNT at ages P7, P14 and P30. P7 (CNT n=7, SEM±6.45; cKO n=6, mean=44.97%, SEM ±5.82, p=0.00006; df=11.00). P14 (CNT n=7, SEM±9.12, cKO n=6, mean=42.87%, SEM±7.37; p=0.0005, df=10.84). P30 (CNT n=8, SEM±2.27; cKO n=5, mean=61.25%, SEM±10.16; p=0.0172, df=4.403). **(C) pPKCα/β in corpus callosum compared to spinal cord at ages P7, P14 and P30**. (D) total PCKα at ages P7, P14 and P30. P7 (CNT n=7, SEM±5.547; cKO n=6, mean=51.30%, SEM±6.691, p=0.0002; df=10.21). P14 (CNT n=7, SEM±11.41, cKO n=6, mean=61.84, SEM ±9.282; p=0.0252, df=10.86). P30 (CNT n=8, SEM±15.86; cKO n=5, mean=50.10%, SEM±12.43; p=0.0308, df=10.99). (E) pS6RP at P14 and P30. P14 (CNT n=7, SEM±9.22, cKO n=6, mean=86.88%, SEM±16.86; p=0.5142, df=7.849). P30 (CNT n=8, SEM±4.94; cKO n=5, mean=102.4%, SEM±15.61; p=0.8900, df=4.812). **(F) p-mTOR-S2448 at P14 and P30**. P14 (CNT n=7, SEM±14.15, cKO n=6, mean=112.6%, SEM±11.91; p=0.5102, df=10.93). P30 (CNT n=8, SEM±6.29; cKO n=5, mean=83.84%, SEM±20.95; p=0.4952, df=4.732). **(G) pAKT-T308 at P14 and P30**. P14 (CNT n=7, SEM±10.21, cKO n=6, mean=125.5%, SEM±8.99; p=0.0878, df=10.99). P30 (CNT n=8, SEM±7.52; cKO n=5, mean=94.85, SEM±7.10; p=0.6286, df=10.48). CNT mean set to 100% based on averaging of actual numerical values. Values are displayed as ±SEM. Unpaired t-test with Welch’s correction.

### Impact of Rictor loss in developing oligodendrocytes on the number of myelinated axons

Ablation of mTORC2 signaling in OPCs reduced differentiation of OPCs to CC1+ maturing oligodendrocytes and overall myelin production in corpus callosum to a much greater extent than in spinal cord. We therefore studied how myelination of individual axons was impacted in corpus callosum and spinal cord by electron microscopy at P21 and 1.5 months (1.5M) (Figure 7). The density of myelinated axons in Rictor cKO corpus callosum genu at P21 was not significantly reduced; however, at 1.5M the density of myelinated axons was significantly reduced. The density of unmyelinated axons was increased at 1.5M but given high variability among the Rictor cKO samples, the increase was not statistically significant (p value=0.0528) (Figure 7A,B). Importantly, however, when g-ratios, a standard measure of myelin thickness, were quantified at either P21 or 1.5M, the g-ratios for Rictor cKO or control corpus callosum myelinated axons were comparable, indicating that while fewer axons were myelinated, those that were myelinated had normal myelin thickness (Figure 7C). The relative frequency distribution of the diameter of myelinated axons shows a slight shift to thinner diameter axons in the Rictor cKO corpus callosum compared to control for both P21 and 1.5M.

By contrast, no significant change in the density of myelinated or unmyelinated axons in Rictor cKO spinal cord was noted at either P21 or 1.5M, relative to controls (Figure 7D,E). As with Rictor cKO corpus callosum, there was no significant change in g ratios of myelinated axons in the spinal cord of Rictor cKO mice relative to control (Figure 7F). The relative frequency distribution of axon diameters in the spinal cord showed no difference in the Rictor cKO compared to control at P21 and 1.5M.

**Figure 7:**
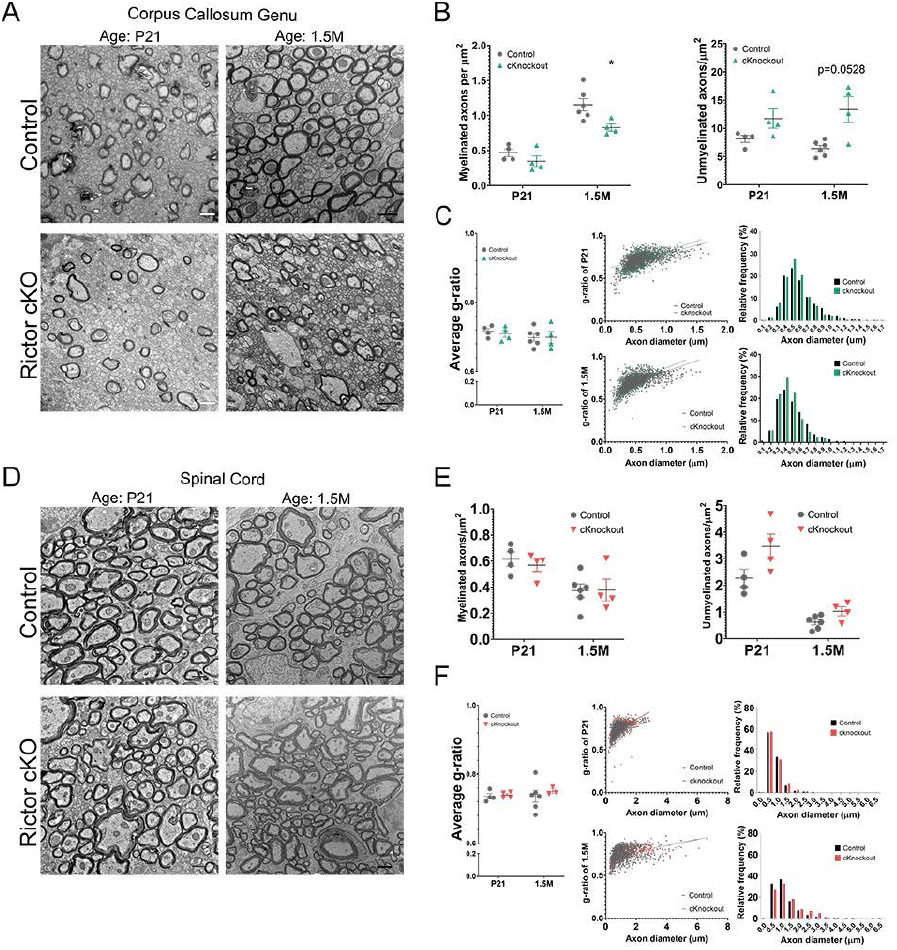
Loss of Rictor results in regional differences in the number of myelinated axons and unmyelinated axons when analyzing the corpus callosum, spinal cord and optic nerve. (**A**) Transmission electron microscopy (TEM) images of the corpus callosum (genu) at 1.5 months in Rictor cKO (cKO) compared to control (CNT). (**B**) Quantification of myelinated and unmyelinated axons of TEM images. Left: P21 myelinated axons per um^2^ (CNT n=4, SEM±0.054, mean=0.4667 axons per um^2^; cKO n=4, SEM±0.077; mean=0.3479 axons per um^2^, p=0.2589, df=5.348); 1.5-M-myelinated axons per um^2^ (CNT n=6, SEM±0.088, mean=1.155 axons per um^2^; cKO n=4, SEM ±0.0504; mean=0.8287 axons per um^2^, p=0.0135, df=7.174). Right: P21 unmyelinated axons per um^2^ (CNT n=4, SEM±0.6237, mean=8.138 axons per um^2^; cKO n=4, SEM±1.707; mean=11.74 axons per um^2^, p=0.1227, df=3.787); 1.5M unmyelinated axons per um^2^ (CNT n=6, SEM±0.5392, mean=6.434 axons per um^2^; cKO n=4, SEM±2.276; mean=13.32 axons per um^2^, p=0.0528, df=3.34). (**C**) Quantification of g-ratio of myelinated axons in TEM images. (Left) P21 Average g-ratio (CNT n=4, SEM±0.0082, mean=0.7165 g-ratio; cKO n=4, SEM±0.0099; mean=0.7088 g-ratio, p=0.5737, df=5.789); 1.5M-Average g-ratio (CNT n=6, SEM±0.0105, mean=0.6982 g-ratio; cKO n=4, SEM±0.0172; mean=0.6989 g-ratio, p=0.9767, df=5.223). (Middle) Distribution of g-ratio by axon diameter (µm). P21 Simple linear regression (CNT Slope=0.1855, y-intercept=0.6097, Slope SEM=0.0069, R^2^=0.3115; cKO Slope=0.2471, y-intercept=0.5662, Slope SEM=0.0099, R^2^=0.3454) 1.5M (CNT Slope=0.2495, y-intercept=0.5768, Slope SEM=0.0067, R^2^=0.4085; cKO Slope=0.2996, y-intercept=0.5652, Slope SEM=0.0119, R^2^=0.3819). (Right) Relative frequency percent of myelinated axon diameters for P21 and 1.5M in the corpus callosum for control and Rictor cKO mice, bin 0.5µm. (**D**) TEM images of the spinal cord in Rictor cKO compared to control at ages P21 and 1.5M. (**E**) Quantification of myelinated and unmyelinated axons of EM images. (Left) P21 myelinated axons per um^2^ (CNT n=4, SEM±0.0555, mean=0.6157 axons per um^2^; cKO n=4, SEM±0.0480; mean=0.5685 axons per um^2^, p=0.5446, df=5.878) 1.5M myelinated axons per um^2^ (CNT n=6, SEM±0.0506, mean=0.3733 axons per um^2^; cKO n=4, SEM±0.0836; mean=0.3783 axons per um^2^, p=0.9618, df=5.184). (Right) P21 unmyelinated axons per um^2^ (CNT n=4, SEM±0.3308, mean=2.279 axons per um^2^; cKO n=4, SEM±0.4714; mean=3.462 axons per um^2^, p=0.0912, df=5.378); 1.5M unmyelinated axons per um^2^ (CNT n=6, SEM±0.102, mean=0.6224 axons per um^2^; cKO n=4, SEM±0.1711; mean=1.03 axons per um^2^, p=0.0946, df=5.125). (**F**) Quantification of g-ratio of myelinated axons in EM images. (Left) P21 Average g-ratio (CNT n=4, SEM ±0.0077, mean=0.7359 g-ratio; cKO n=4, SEM ±0.0044; mean=0.7398 g-ratio, p=0.6761, df=4.776); 1.5M-Average g-ratio (CNT n=6, SEM ±0.0169, mean=0.7361 g-ratio; cKO n=3, SEM ±0.0062; mean=0.751 g-ratio, p=0.4383, df=6.159). (Middle) Distribution of g-ratio by axon diameter (µm). P21 Simple linear regression (CNT Slope=0.0976, y-intercept=0.6595, Slope SEM±0.0035, R^2^=0.3242; cKO Slope=0.0993, y-intercept=0.6607, Slope SEM=0.0038, R^2^=0.3377) 1.5M (CNT Slope=0.03559, y-intercept=0.7084, Slope SEM±0.0029, R^2^=0.1114; cKO Slope=0.0370, y-intercept=0.7030, Slope SEM=0.0028, R^2^=0.236). (Right) Relative frequency percent of myelinated axon diameters for P21 and 1.5M in the spinal cord for control and Rictor cKO mice. Values are displayed as ±SEM. Unpaired t-test with Welch’s correction or simple linear regression. TEM images scale bar 1µm.

### Rictor conditional deletion results in altered actin cytoskeleton regulators

An important downstream target of mTORC2 signaling is cytoskeletal regulation, specifically actin. Several models of active myelination indicate that the cycling of F-actin and G-actin is involved in the production of myelin (Nawaz et al., 2015; Zuchero et al., 2015). An *in vivo* stain allows quantification of the F-actin/G-actin ratio in tissue, and consistent with studies by Nawaz et al (2015), the F-actin/G-actin ratio is higher in gray matter areas than white matter areas (Figure 8A-D), in cortex relative to corpus callosum at both P14 and P30. However, that ratio was not altered in the Rictor cKO mice compared to control. On the other hand, it was clear that the overall width of the corpus callosum was significantly reduced in the Rictor cKO compared to control at P14 and P30 (Figure 8E). This likely resulted from the reduction in myelinated axons in corpus callosum of Rictor cKO mice (Figure 7).

**Figure 8:**
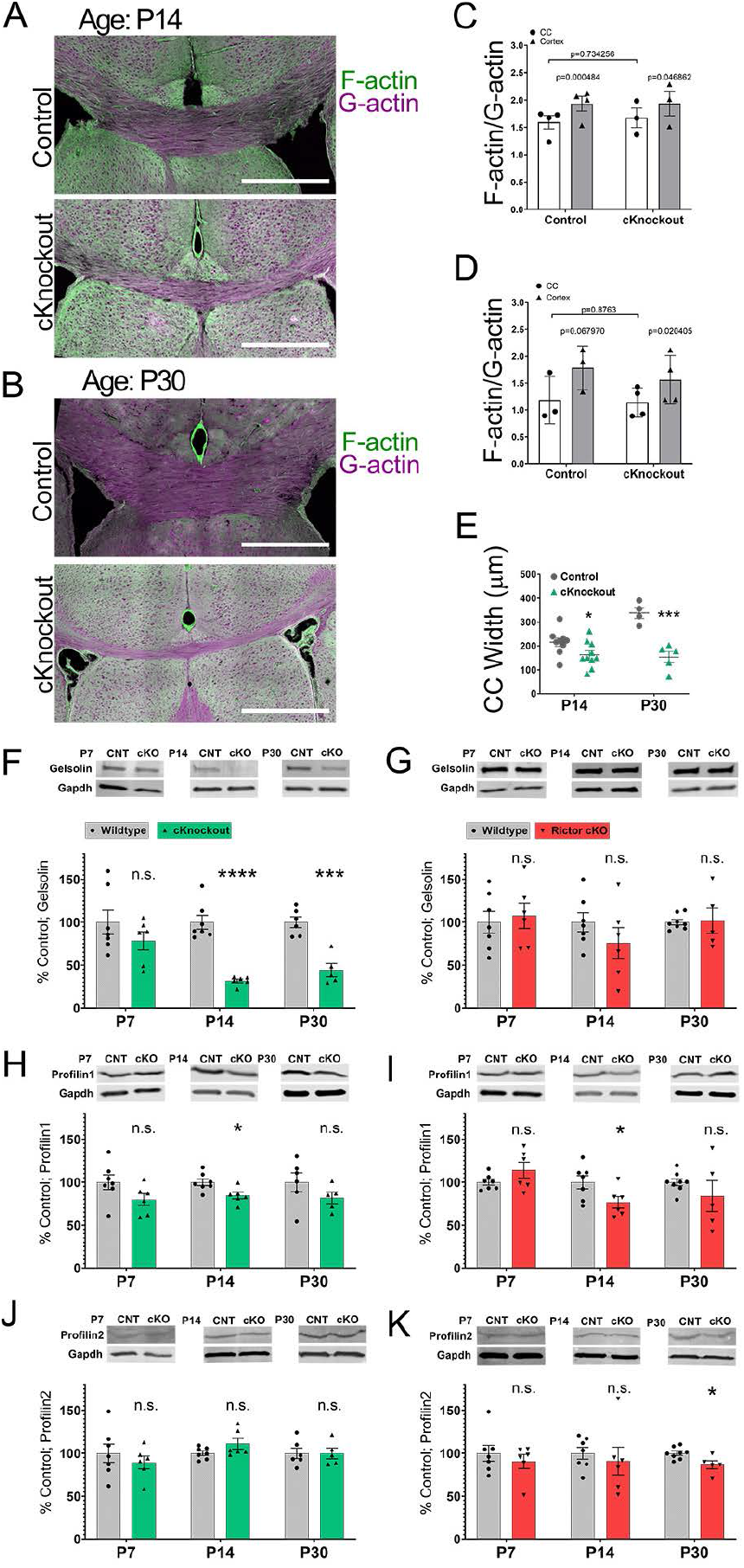
Loss of mTORC2 signaling has little impact on G-actin/F-actin ratio in corpus callosum but results in decreased Gelsolin. (**A**) Corpus callosum of P14 animals stained for F-actin (Green, phalloidin) and G-actin (Magenta, DNAseI) comparing Rictor cKO with control (scale bar 500µm). (**B**) Corpus callosum of P30 animals stained for F-actin (Green) and G-actin (Magenta) comparing Rictor cKO with control (scale bar 500µm). (**C**) Ratio of F-actin to G-actin intensity at age P14 in white matter (open bars) or gray matter (gray bars). Control (n=4, SEM±0.122, mean=1.596, paired t-test p=0.00005, df=4.0) cKO (n=3, SEM±0.178, mean=1.675, paired t-test p= 0.0469, df=3.0); Control CC vs cKO CC (p= 0.3212, df=3.736). (**D**) Ratio of F-actin to G-actin intensity at age P30 in white matter (open bars) or gray matter (gray bars). Control (n=3, SEM±0.256, mean=1.186, paired t-test p=0.0680, df=2.0) cKO (n=4, SEM±0.137, mean=1.137, paired t-test p= 0.0204, df=3.0); Control CC vs cKO CC (p= 0.8763, df=3.135). (**E**) Corpus callosum widths were measured using F-actin/G-actin stained images at the approximate midline of control compared to Rictor cKO mice in P14 and P30. P14 (CNT n=9, SEM±17.249, mean=215.778 µm, cKO n=10, mean=163.444 μm, SEM±17.059, p=0.0457, df=16.92) P30(CNT n=4, SEM±22.614, mean=338.300μm, cKO n=5, SEM±23.702, mean=153.538 µm, p=0.0008, df=6.935); Control P14 vs P30 (p=0.0040, df=6.661) cKO P14 vs P30 (p=0.7429, df=8.235). **(F-K)**Quantification of actin regulators in corpus callosum of CNT and cKO mice at P7, P14 and P30. (**F**) Western blot (WB): **Gelsolin** in corpus callosum; CNT and cKO at P7, P14 and P30. Gelsolin-P7 (CNT n=7, SEM±13.98; cKO n=6, SEM±10.63; cKO mean=78.35%, p=0.2441, df=10.66), P14 (CNT n=7, SEM±8.03; cKO n=6, SEM±2.171; cKO mean=31.65%, p=0.0001, df=6.865), P30 (CNT n=6, SEM±6.32; cKO n=5, SEM±7.715; cKO mean=44.11%, p=0.0005, df=8.212). (**G**) WB: **Gelsolin** in the spinal cord at ages P7, P14 and P30. P7 (CNT n=7, SEM±12.5; cKO n=6, SEM±14.79; cKO mean=107.5%, p=0.7057, df=10.31), P14 (CNT n=7, SEM±11.01; cKO n=6, SEM±17.96; cKO mean=75.8%, p=0.2821, df=8.468), P30 (CNT n=8, SEM±3.07; cKO n=5, SEM±14.78; cKO mean=101.5%, p=0.9228, df=4.348). (**H**) WB: **Profilin1** in corpus callosum at ages P7, P14 and P30. Profilin1-P7 (CNT n=7, SEM±8.497; cKO n=6, SEM±6.915; cKO mean=79.72%, p=0.0915, df=10.86), P14 (CNT n=7, SEM±3.94; cKO n=6, SEM±3.79; cKO mean=84.39%, p= 0.0157, df= 10.97), P30 (CNT n=6, SEM±10.83; cKO n=5, SEM±6.58; cKO mean=81.59%, p=0.1843, df=8.01). (**I**) WB: **Profilin1** in the spinal cord at ages P7, P14 and P30. Profilin1 P7 (CNT n=7, SEM±3.33; cKO n=6, SEM±9.19; cKO mean=114.0%, p= 0.1996, df= 6.31), P14 (CNT n=7, SEM±7.60; cKO n=6, SEM±6.76; cKO mean=76.52%, p=0.0414, df=10.99), P30 (CNT n=8, SEM±4.10; cKO n=5, SEM±17.99; cKO mean=83.83%, p=0.4260, df=4.42). (**J**) WB: **Profilin2** P7 (CNT n=7, SEM±10.62; cKO n=6, SEM±7.44; cKO mean=89.32%, p=0.4289, df=10.34), P14 (CNT n=7, SEM±2.56; cKO n=6, SEM±6.27; cKO mean=111.0%, p=0.1503, df=6.65), P30 (CNT n=6, SEM±6.03; cKO n=5, SEM±6.26; cKO mean=99.85%, p= 0.9870, df= 8.80). (**K**) WB: **Profilin2** P7 (CNT n=7, SEM±9.14; cKO n=6, SEM±8.18; cKO mean=90.50%, p=0.4552, df=11.00), P14 (CNT n=7, SEM±6.86; cKO n=6, SEM±16.12; cKO mean=90.97%, p=0.6225, df=6.79), P30 (CNT n=8, SEM±2.41; cKO n=5, SEM±4.70; cKO mean=86.58%, p=0.0432, df=6.14).

Despite the comparable ratio of F-actin/G-actin, there were important differences in some regulators of actin polymerization in the Rictor cKO mice. When quantified by western blot, profilin1 was somewhat reduced through development in Rictor cKO corpus callosum, while profilin2 was unchanged, and these actin regulators were only minimally reduced in Rictor cKO spinal cord (Figure 8H-K). Importantly, gelsolin was dramatically reduced at P14 and P30 in Rictor cKO corpus callosum, while it was not significantly affected in the spinal cord of Rictor cKO mice at any age (Figure 8F-G). These data would suggest that gelsolin regulation of actin depolymerization may be a downstream effector of mTORC2 signaling regulating the cytoskeleton, particularly in corpus callosum, but not in spinal cord, where it remains normally expressed and myelination occurs normally, despite loss of mTORC2 signaling.

## Discussion

The production of myelin by oligodendrocytes is complex and requires precise temporal and spatial regulation of intracellular signals that control the various activities required for this crucial developmental process. While previous studies have shown that mTOR is important for oligodendrocyte differentiation and myelin formation, the effectors of mTOR signaling that drive differentiation and myelin formation are still not fully understood.

In earlier studies, we established that the impact of mTOR signaling in oligodendrocytes is different, depending on the CNS region that is examined (Bercury et al., 2014; Wahl et al., 2014). Importantly, in the earlier studies, we established the loss of mTOR itself or of Raptor (mTORC1) resulted in reduced myelination in spinal cord, but not in corpus callosum. In those studies, loss of Rictor (mTORC2) had little impact in either region (Bercury et al., 2014). In those studies and a comparable study from the Suter laboratory (Lebrun-Julien et al., 2014), mTOR, Raptor or Rictor were deleted from oligodendrocyte lineage cells using the CNP promoter. CNP is typically considered a late marker of oligodendrocyte development; by single cell RNAseq analysis (Marques et al., 2016), CNP RNA is barely detectable in OPCs. Those studies clearly showed that deletion of mTORC2 signaling from late differentiating/myelinating oligodendrocytes had relatively little impact on myelination throughout the CNS.

By contrast, in the current study we establish that mTORC2 does indeed have a role in oligodendrocyte development, and it is detrimental when it is deleted early in OPC differentiation. An earlier study by Grier et al (2017) established that Rictor conditional deletion from all Olig2-positive cells resulted in reduced oligodendrocytes and myelination in cortex, and they suggested that the reduction in oligodendrocyte number might result from an embryonic impact of Rictor loss in Olig2-positive cells, presumably from very early progenitors. The current studies confirm the results of Grier et al (2017) on the impact of Rictor deletion but would not support the conclusion that this outcome results from an embryonic impact on oligodendrocyte cell number. Our studies delete Rictor only from PDGFRα-expressing cells, which would not be embryonic progenitor cells impacted in the earlier Olig2-based model. Nevertheless, Rictor deletion from postnatal OPCs in corpus callosum does result in reduced total oligodendrocytes, with a far greater impact on the number of maturing oligodendrocytes (Figure 2).

More importantly, the current studies extend the analysis of the impact of mTORC2 on oligodendrocyte development to spinal cord oligodendrocytes. Intriguingly, there was also a different regional impact of mTORC2 loss, but it was unexpected relative to the earlier studies on mTOR or Raptor loss (Bercury et al., 2014; Lebrun-Julien et al., 2014; Wahl et al., 2014). Thus, in the Rictor cKO mice investigated here, differentiation and myelination in spinal cord was relatively spared, but its loss from OPCs specifically in the corpus callosum had a dramatic impact reducing myelination. Our main finding was that OPC-specific knockout of Rictor and loss of mTORC2 signaling resulted in reduced numbers of oligodendrocytes, reduced differentiation of those oligodendrocytes and fewer myelinated axons in corpus callosum in Rictor cKO animals compared to control, but not in spinal cord, the exact opposite regional impact of Raptor or mTOR loss from more mature oligodendrocytes (Bercury et al., 2014; Lebrun-Julien et al., 2014; Wahl et al., 2014).

The current studies focus extensively on the signaling impact of the loss of mTORC2 in OPCs. Importantly, the impact of Rictor loss on oligodendrocyte differentiation during early brain development in corpus callosum did not alter mTORC1 signaling (Figure 4). However, by P30 the loss of mTORC2 in later stages of corpus callosum myelination did alter mTORC1 signaling, reducing S6RP phosphorylation. Interestingly, neither pAkt-T308, the Akt phosphorylation substrate of phosphoinositide-dependent kinase 1 (PDK1) driving mTORC1 signaling, nor phosphorylation of mTOR itself at S2448, which is the S6RP negative feedback regulator of mTORC1 signaling (Figueiredo, 2017), were affected at late stages of development. Likely the overall impact on myelination resulting from loss of mTORC2 signaling reduced oligodendrocyte differentiation in corpus callosum in general.

The regional differences in mTORC2 requirement seen in this study did not result from an inability to delete Rictor or to impact the known function of mTORC2 to phosphorylate Akt at S473, as spinal cord clearly had reduced Rictor and p-AktS473 (Figure 6). However, it may result from differences in endogenous PKCα or PKCβ_II_ signaling, as we found that endogenous phosphorylation of PKCα/β_II_ was not detected in the spinal cord from P7 to P30 (Figure 6). Since phospho-PKCα/β_II_ was present in control corpus callosum and it was significantly downregulated in Rictor cKO corpus callosum, this may be a major driver of mTORC2 signaling during OPC differentiation in corpus callosum, quite distinct from OPC differentiation in spinal cord.

One of our recent studies (Musah et al., 2020) showed that the inhibition of mTOR signaling results in a decrease in Profilin2, a member of the profilin family that is quite abundant in the brain. However, deleting mTORC2 signaling in OPCs in the current study resulted in no change in Profilin2 in corpus callosum. Thus, mTORC1 is likely responsible for the loss of Profilin2 in mTOR cKO mice, where it has a greater impact on spinal cord myelination. On the other hand, loss of mTORC2 did result in loss of the more ubiquitous Profilin1 at P14, which may indicate some direct regulation of Profilin1 expression by mTORC2. However, Profilin1 was reduced in both corpus callosum and spinal cord at P14, suggesting that it does not drive the differential impact of Rictor loss from OPCs in corpus callosum and spinal cord. Our prior study (Musah et al., 2020) also indicated loss of mTOR regulates a variety of cytoskeletal processes in differentiating oligodendrocytes in spinal cord. The lack of a major effect of Rictor loss on oligodendrocyte differentiation in the spinal cord suggests much of this occurs independently of mTORC2, likely through mTORC1, whereas the corpus callosum relies more on mTORC2 for oligodendrocyte differentiation. This is also supported by our prior findings on the Raptor conditional knockout mice, where oligodendrocyte differentiation was compromised specifically in spinal cord (Bercury et al., 2014).

More importantly, the loss of mTORC2 signaling resulted in a dramatic loss of gelsolin in corpus callosum from P14, i.e., the time of active OPC differentiation and myelination. No loss of gelsolin was seen in spinal cord, which could be a key difference in the impact of mTORC2 deletion from OPCs. Gelsolin is a Ca2+- and polyphosphoinositide 4,5-bisphosphate (PIP2)-regulated actin binding protein whose major activity is in filamentous actin severing, which it does with close to 100% efficiency (Selden et al., 1998). Gelsolin is involved in oligodendrocyte differentiation and myelination. It is highly expressed in the oligodendrocyte lineage, with greatest expression in actively myelinating cells, but with significant expression in newly formed oligodendrocytes (Zhang et al., 2014). It is proposed to be sequestered at the plasma membrane prior to the production of MBP, which then displaces it at the membrane, releasing this actin-severing protein to drive actin disassembly (Zuchero et al., 2015). In its absence, the actin polymerization/depolymerization cycle would be nonfunctional, reducing the ability to generate compact myelin.

The current studies establish that in the corpus callosum OPC differentiation is regulated by mTORC2 signaling. However, while there were fewer myelinated axons in Rictor cKO corpus callosum, among the myelinated axons in corpus callosum, there was little difference in g-ratio in Rictor cKO corpus callosum relative to controls. This indicates that, while fewer OPCs in the Rictor cKO mice were able to differentiate, those that did progress to myelination ensheathed axons with the appropriate amount of myelin. It was noted that a larger proportion of myelinated axons in the Rictor cKO were thinner than in control. This suggests that the smaller axons in the Rictor cKO are more likely to be myelinated. Interestingly, in the Grieg et al. (2017) study, myelinated axon diameter in the cKO corpus callosum was thicker than control, suggesting that deletion of Rictor at different stages of OPC development may bias axon selection. These studies suggest that mTORC2 signaling appears important for the initial differentiation of OPCs and premyelinating cells, but once cells differentiated beyond the earliest stages of myelination, other regulators drive the late stages of myelination. This may explain the relatively minor impact of mTORC2 loss in CNP-Cre mice, where the deletion would have been at the later stages of oligodendrocyte differentiation. This may also explain the recovery seen in the Olig2-Cre-deleted mice as they age (Grier et al., 2017). Thus, it may be in that model, sufficient cells are able to escape loss of mTORC2 signaling during differentiation and once past that stage, they are able to produce essentially normal myelin. Unfortunately, because of significant PDGFRα promoter-driven gene expression in embryonic heart (Klinghoffer et al., 2001), the PDGFRα-Cre mice in the present studies likely deleted mTORC2 signaling in developing heart and these mice did not survive significantly past 2 months of age.

Further examination of the role of mTORC2 in oligodendrocyte development is necessary to elucidate the full impact of this kinase. We still do not know the direct targets of mTORC2 in oligodendrocytes. It is intriguing to speculate about the relative importance of mTORC2 signaling at different stages of oligodendrocyte differentiation. Clearly both this study and Grier et al.(2017) demonstrate that deletion of Rictor is detrimental to OPC differentiation in corpus callosum, while our earlier study with CNP-Cre driven deletion of Rictor (Bercury et al., 2014) showed normal oligodendrocyte differentiation but early loss of myelin RNAs from corpus callosum with some recovery by P29, and essentially normal myelin thickness by 2 months of age. Thus, loss of mTORC2 signaling after early OPC differentiation impacted gene expression, but there was sufficient recovery that normal myelin was produced. Both our study (Bercury et al., 2014) and that of Lebrun-Julien et al. (2014) showed little impact in spinal cord, but the current study suggests no impact would have been expected irrespective of the timing of Rictor deletion from spinal cord oligodendrocytes. Studies investigating the impact of Rictor/mTORC2 signaling with respect to cytoskeleton reorganization in the differentiating oligodendrocyte are underway to determine how mTORC2 signaling regulates OPC differentiation.

## Acknowledgements

This work was supported by National Institutes of Health R37 #82203 to W.B.M. and T.L.W; by National Science Foundation GRFP DGE-1553798 to T.L.B. and NIH 5F31NS118830 to A.R.A. We thank members of the W.B.M. and T.L.W. laboratories for valuable discussion and feedback on this manuscript.

## Author contributions

H.A.H., K.D.D. and W.B.M. designed research; K.D.D., A.R.A. H.A.H., T.L.B., L.T.F. and J.B. performed research; K.D.D., A.R.A. and W.B.M. analyzed data; K.D.D. wrote the first draft of the paper; all authors edited the paper.

## Notes

### Competing Interest Statement

The authors have declared no competing interest.

